# GPR27 mediates L-lactate-induced Ca^2+^ and cAMP signalling in 3T3 cells

**DOI:** 10.64898/2026.08.09.743761

**Authors:** Danaja Kuhanec, Ena Sanjković, Tomaž M. Zorec, Marko Kreft, Helena Haque Chowdhury, Robert Zorec

**Affiliations:** Laboratory of Neuroendocrinology-Molecular Cell Physiology, Institute of Pathophysiology, Faculty of Medicine, University of Ljubljana, Zaloška 4, Ljubljana, Slovenia; Celica Biomedical, Tehnološki park 24, Ljubljana, Slovenia; Department of Biology, Biotechnical Faculty, University of Ljubljana, Večna pot 111, Ljubljana, Slovenia

**Keywords:** GPR27, SREB1, orphan GPCR, 8535n, L-lactate, intracellular calcium, cAMP, FRET nanosensor, 3T3 MEF cells

## Abstract

GPR27/SREB1 is a highly conserved orphan class A G-protein coupled receptor implicated in insulin production, metabolic regulation, tumour biology, neurodegeneration and L-lactate homeostasis, but its immediate second-messenger signalling remains poorly defined. We used single-cell Förster resonance energy transfer nanosensors to monitor cytosolic Ca^2+^ and cAMP in wild-type 3T3 MEF cells, CRISPR–Cas9 GPR27-knockout cells (GPR27KO) and GPR27-knockout cells transiently re-expressing FLAG-tagged GPR27 (GPR27-rescued). The GPR27 surrogate agonist 8535n (1 µM) increased intracellular Ca^2+^ in wild-type and rescued cells but not in GPR27-knockout cells and produced no significant cAMP response in wild-type cells. Basal Ca^2+^ and cAMP levels were unaffected by GPR27 deletion. Extracellular L-lactate (2 mM) induced a GPR27-dependent increase in Ca^2+^ and cAMP in wild-type and rescued cells, but not in knockout cells, raising the possibility that L-lactate acts as an endogenous ligand or modulator of GPR27.

**Graphical abstract:** 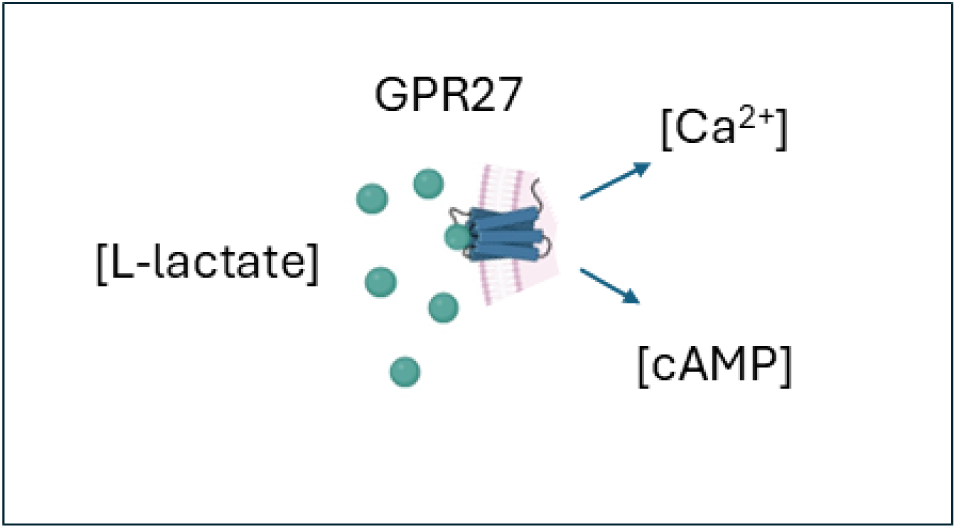

**Highlights:**

- GPR27 surrogate agonist 8535n increases intracellular Ca^2+^ but not cAMP in 3T3 cells.
- Extracellular L-lactate induces GPR27-dependent intracellular Ca^2+^ and cAMP increases in 3T3 cells.
- These findings identify GPR27 as a putative candidate lactate sensor.

## Introduction

G-protein coupled receptors (GPCRs) translate extracellular chemical, mechanical, sensory, and metabolic signals into intracellular responses and represent major therapeutic targets. However, many human GPCRs lack validated endogenous ligands and remain classified as orphan GPCRs (oGPCRs). Characterizing these receptors may reveal previously unrecognized physiological pathways and new therapeutics^1, 2, 3, 4^.

GPR27, also known as SREB1, is a highly conserved class A oGPCR belonging to the Superconserved Receptors Expressed in the Brain family, together with GPR85/SREB2 and GPR173/SREB3^5, 6^. Although prominently expressed in the central nervous system, GPR27 is also found in pancreatic islets, ovaries, testes, prostate and heart, suggesting functions in both neural and peripheral physiology^1, 5, 7^.

GPR27 has been implicated in neurodegeneration^8^ and in several cancers, although its effects appear tumour- and context-dependent. Reduced GPR27 expression has been associated with glioma progression, whereas studies in gastric, ovarian and liver cancers indicate variable relationships with tumour growth, immune infiltration, treatment response and survival^9, 10, 11, 12^. These findings suggest that GPR27 cannot be assigned a universal tumour-promoting or tumour-suppressive role.

More consistent evidence links GPR27 to metabolic regulation. In pancreatic β cells, GPR27 knockdown reduced insulin-promoter activity and glucose-stimulated insulin secretion^13^. GPR27-deficient mice showed reduced islet insulin and Pdx1 expression and lower circulating insulin, although systemic glucose homeostasis was only modestly affected^14^. In zebrafish, *gpr27* deletion aggravated hyperglycaemia, impaired insulin-dependent Akt activation and glucose utilization, and acylcarnitine metabolism^15^.

GPR27 has also been linked to lactate homeostasis. Surrogate GPR27 agonists increased intracellular L-lactate in wild-type (WT) 3T3 murine embryonic fibroblast (MEF) cells and rat astrocytes, whereas this response was reduced by *gpr27* deletion and restored by receptor re-expression. GPR27-knockout cells also displayed increased basal intracellular lactate ([lactate]_i_), supporting a role for GPR27 in cellular lactate regulation^16^. It has been known for some time that lactate homeostasis is altered in cancer cells; many cancer cells produce L-lactate even in the presence of oxygen, a process called aerobic glycolysis^17^.

L-Lactate is both a metabolic substrate and a signalling molecule. Its established receptor, HCAR1/GPR81, typically couples to G_i/o_ and inhibits adenylyl cyclase. In astrocytes, however, extracellular L-lactate and GPR81 agonists increased [lactate]_i_ and cAMP ([cAMP]_i_) also through a GPR81-independent mechanism, suggesting the existence of an additional excitatory lactate-sensing pathway^18, 19, 20, 21^.

The signalling pathways activated by GPR27 remain unresolved. Although several surrogate GPR27 agonists have been developed, the endogenous ligand and physiologically relevant coupling pathways remain unknown^1, 22^. GPR27-dependent phospholipase C/IP_1_ production suggests possible G_q/11_ coupling, whereas other studies have reported constitutive G_i/o_ activity or β-arrestin-2 recruitment without detectable G-protein activation^13, 23, 24, 25^.

Because intracellular Ca^2+^ ([Ca^2+^]_i_) and [cAMP]_i_ responses provide mechanistic information about GPCR coupling, we used genetically encoded Förster resonance energy transfer (FRET) nanosensors to monitor these messengers in individual cells. We compared WT 3T3 cells, CRISPR–Cas9 GPR27-knockout cells (GPR27KO), and knockout cells re-expressing GPR27 (GPR27-rescued), a platform introduced previously^16^. The role of GPR27 has been linked to cancers^9, 10, 11, 12^, which often exhibit increased L-lactate production, a biochemical pathway enabling the availability of biosynthetic intermediates generated by aerobic glycolysis, also known as the Warburg effect^17^. Hence, we tested whether extracellular L-lactate induces GPR27-dependent changes in [Ca^2+^]_i_ or [cAMP]_i_, and whether GPR27 deletion alters their basal levels. We also used a surrogate GPR27 agonist to assess GPR27’s involvement in activating these second-messenger pathways.

The results revealed that the surrogate agonist 8535n increased [Ca^2+^]_i_ in WT and rescued cells, but not in GPR27-knockout cells, and produced no significant response in [cAMP]_i_ in WT cells. Interestingly, enhanced vehicle-induced cAMP responses in knockout cells suggest that GPR27 modulates responses to solution exchange or mechanical stimulation. The addition of extracellular L-lactate, which previously was shown to elevate [cAMP]_i_ in astrocytes from an animal model of intellectual disability, elicited increases in [Ca^2+^]_i_ and [cAMP]_i_, which required the presence of GPR27, indicating the possibility that L-lactate acts as an endogenous ligand or modulator of GPR27.

## Results

### The GPR27 surrogate agonist 8535n increases [Ca^2+^]_i_ but not [cAMP]_i_ in 3T3 cells

First, we quantified how GPR27 activation affects the second messengers Ca^2+^ and cAMP using genetically encoded FRET nanosensors. The D3cpv nanosensor^25^ was used for [Ca^2+^]_i_ measurements (Fig. 1).

**Fig. 1.**
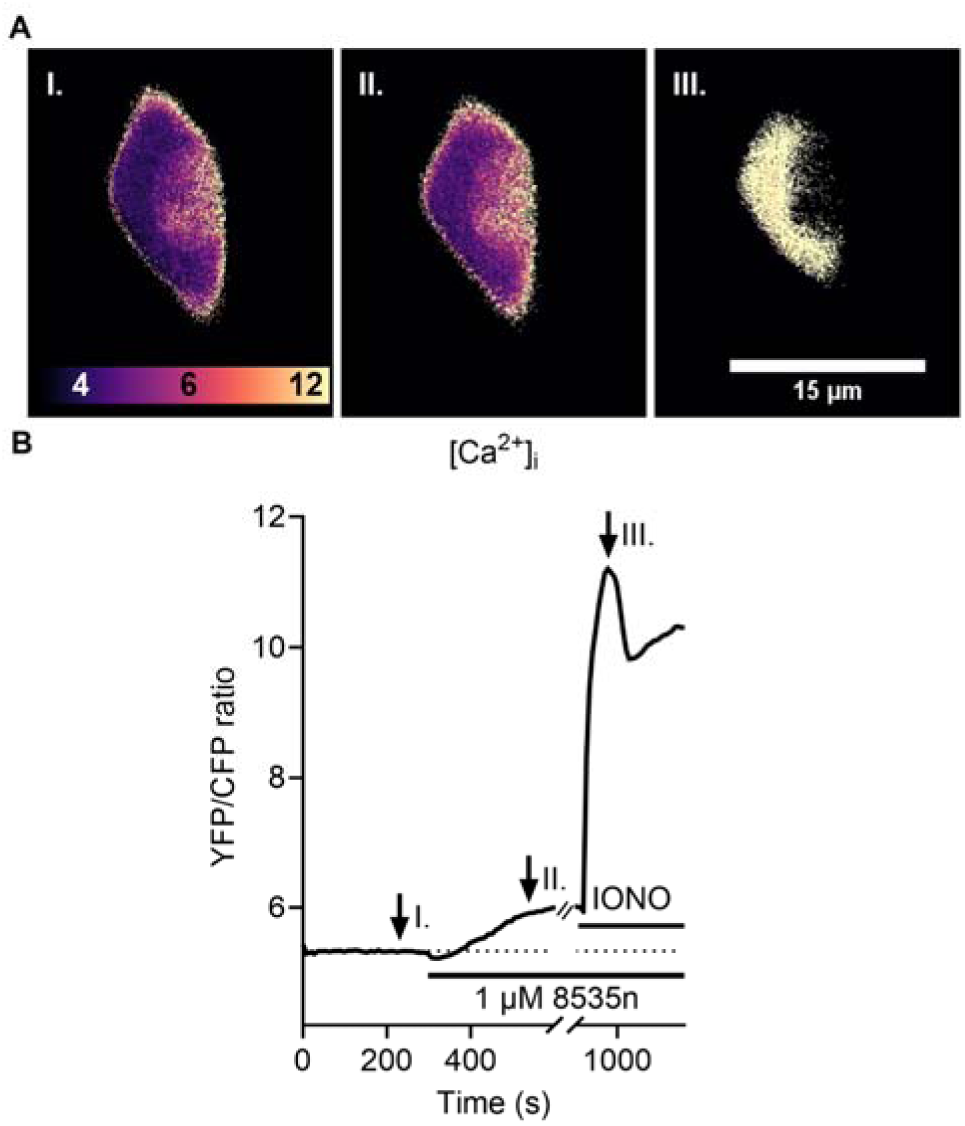
Representative single-cell [Ca^2+^]_i_ response to 8535n and ionomycin in a 3T3 WT cell. (**A**) Pseudocoloured micrographs showing the YFP/CFP ratio before stimulation (I), after 1 µM 8535n (II) and after 10 µM ionomycin (IONO; III) in a single 3T3 WT cell. Scale bar: 15 µm. (**B**) YFP/CFP ratio from the same cytoplasmic region over time. Note the rise in the signal after application of 1 µM 8535n and the larger response to ionomycin, an agent that increases the membrane permeability for Ca^2+^. The dotted horizontal line indicates the baseline YFP/CFP ratio; horizontal bars indicate 8535n and IONO application. Arrows indicate the time points shown in (**A**). For clarity, every third data point is shown. CFP, cyan fluorescent protein; YFP, yellow fluorescent protein.

We used the surrogate agonist 8535n (1 µM) (N-[4-(anilinosulphonyl)phenyl]-2,4-dichlorobenzamide)^1^ to stimulate WT, GPR27KO and rescued cells (GPR27KO + pFLAG27), and vehicle controls received an equal volume of extracellular solution (ECS) to match solution exchange and potential mechanical perturbation. At the end of the experiment, ionomycin (10 µM), a calcium-selective ionophore that increases membrane permeability to Ca^2+^ and increases intracellular Ca^2+^ concentrations^26^, was added, serving as a positive control.

For clarity, FRET recordings are shown from 60 to 600 s, with every third data point plotted (Fig. 2). Responses were quantified as the area under the curve (AUC) relative to baseline during the 300 s after stimulation. Mean [Ca^2+^]_i_ AUC ± SEM values (% × s) were: WT vehicle, 5.17 ± 0.94 (*n* = 12); WT 8535n, 13.02 ± 2.49 (*n* = 12); GPR27KO vehicle, 5.52 ± 0.92 (*n* = 13); GPR27KO 8535n, 3.54 ± 0.90 (*n* = 11); GPR27KO + pFLAG27 vehicle, 8.45 ± 1.30 (*n* = 10); and GPR27KO + pFLAG27 8535n, 15.47 ± 1.47 (*n* = 12). The 8535n response was significant in WT cells (Mann–Whitney test, \**P ≤* 0.05, *U* = 28) and GPR27-rescued cells (Student’s *t* test, \*\**P ≤* 0.01, *t* = −3.5), but not in GPR27KO cells (*P* = 0.141, *t* = 1.5). Vehicle responses did not differ significantly among the groups (one-way analysis of variance [ANOVA], *P* = 0.078, *F* = 2.8), whereas 8535n responses differed significantly (one-way ANOVA with Holm-Šidák post hoc testing, ^###^*P ≤* 0.001, *F* = 12.1; *t*_WT vs. KO_ = 3.7, *t*_WT vs. KO + p27_ = 1.0, t_KO + p27 vs. KO_ = 4.7). Thus, 8535n increased [Ca^2+^]_i_ in WT and GPR27-rescued cells, but not in GPR27KO cells, supporting a GPR27-dependent Ca^2+^ response.

**Fig. 2.**
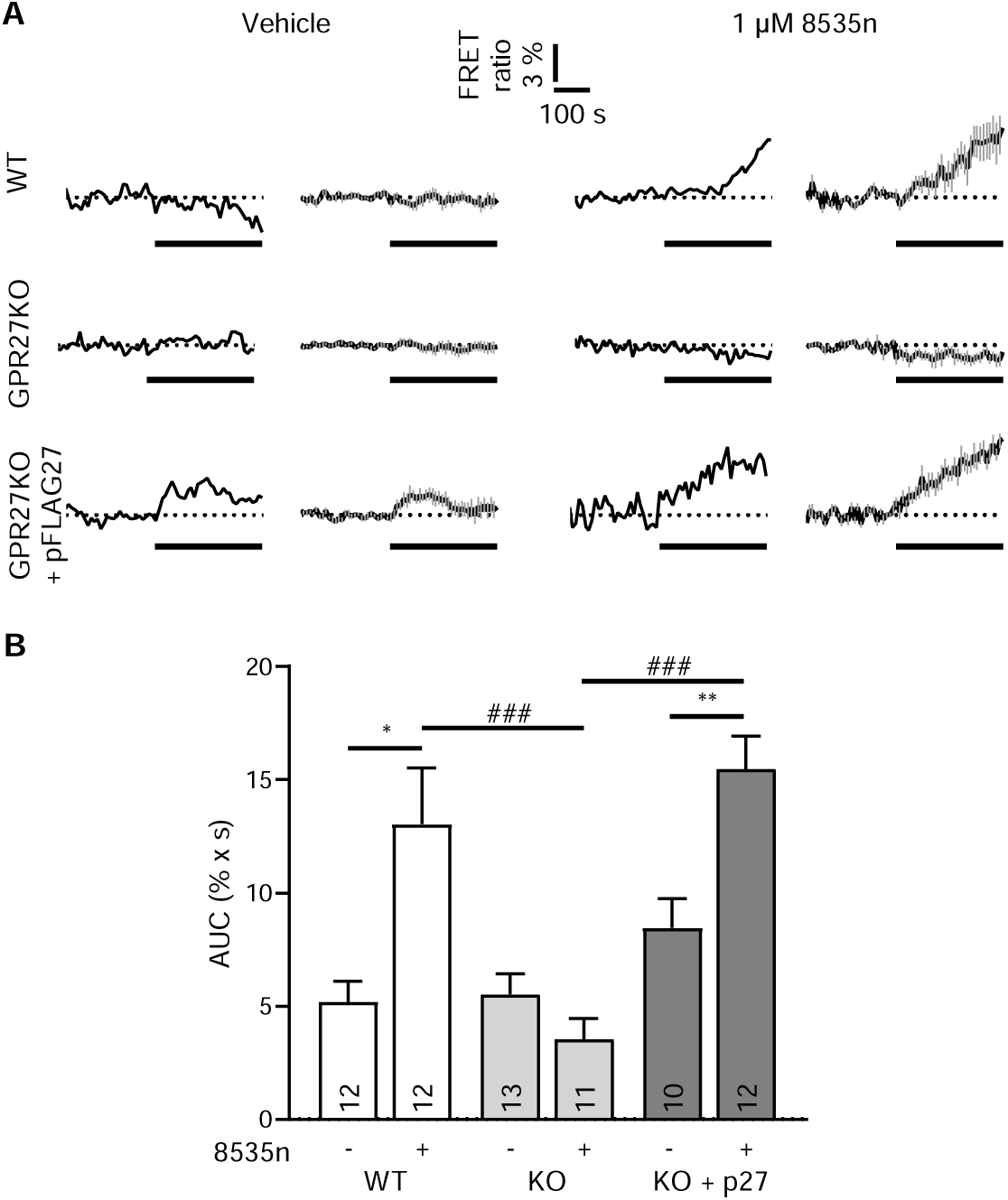
The GPR27 surrogate agonist 8535n increases [Ca^2+^]_i_ in 3T3 WT cells, and the response is rescued by re-expression of GPR27 in GPR27KO cells. (**A**) Representative traces (first and third columns) and averaged traces (mean ± SEM; second and fourth columns) of the normalized YFP/CFP FRET ratio in 3T3 WT, 3T3 GPR27KO and GPR27KO cells transiently re-expressing FLAG-tagged GPR27 (GPR27KO + p27). Cells were treated with vehicle (ECS; left) or 1 µM 8535n (right). Dotted horizontal lines indicate the normalized baseline (0%), and bold horizontal bars indicate stimulus application. FRET recordings are plotted from 60 to 600 s, with every third point shown. (**B**) Mean AUC ± SEM of the normalized [Ca^2+^]_i_ response after vehicle or 8535n. Numbers in the bars indicate independently recorded single-cell observations from at least three cell passages. Statistical significance was assessed by Mann–Whitney rank-sum test for WT (\**P* ≤ 0.05), Student’s *t* test for GPR27KO (*P* = 0.141) and GPR27KO + p27 (\*\**P* ≤ 0.01), and one-way ANOVA followed by Holm-Šidák post hoc test for vehicle (*P* = 0.078) and 8535n-treated groups (^###^*P* ≤ 0.001). AUC, area under the curve; CFP, cyan fluorescent protein; ECS, extracellular solution; FRET, Förster resonance energy transfer; KO, knockout; WT, wild-type; YFP, yellow fluorescent protein.

The experiment (Fig. 3) measuring [cAMP]_i_. with the FRET nanosensor Epac1-camps^27^ shows an increase in cAMP on the application of extracellular L-lactate, followed by the addition of noradrenaline (NA), which is known to increase [cAMP]_i_ and serves as a positive control, as reported previously^28^.

**Fig. 3.**
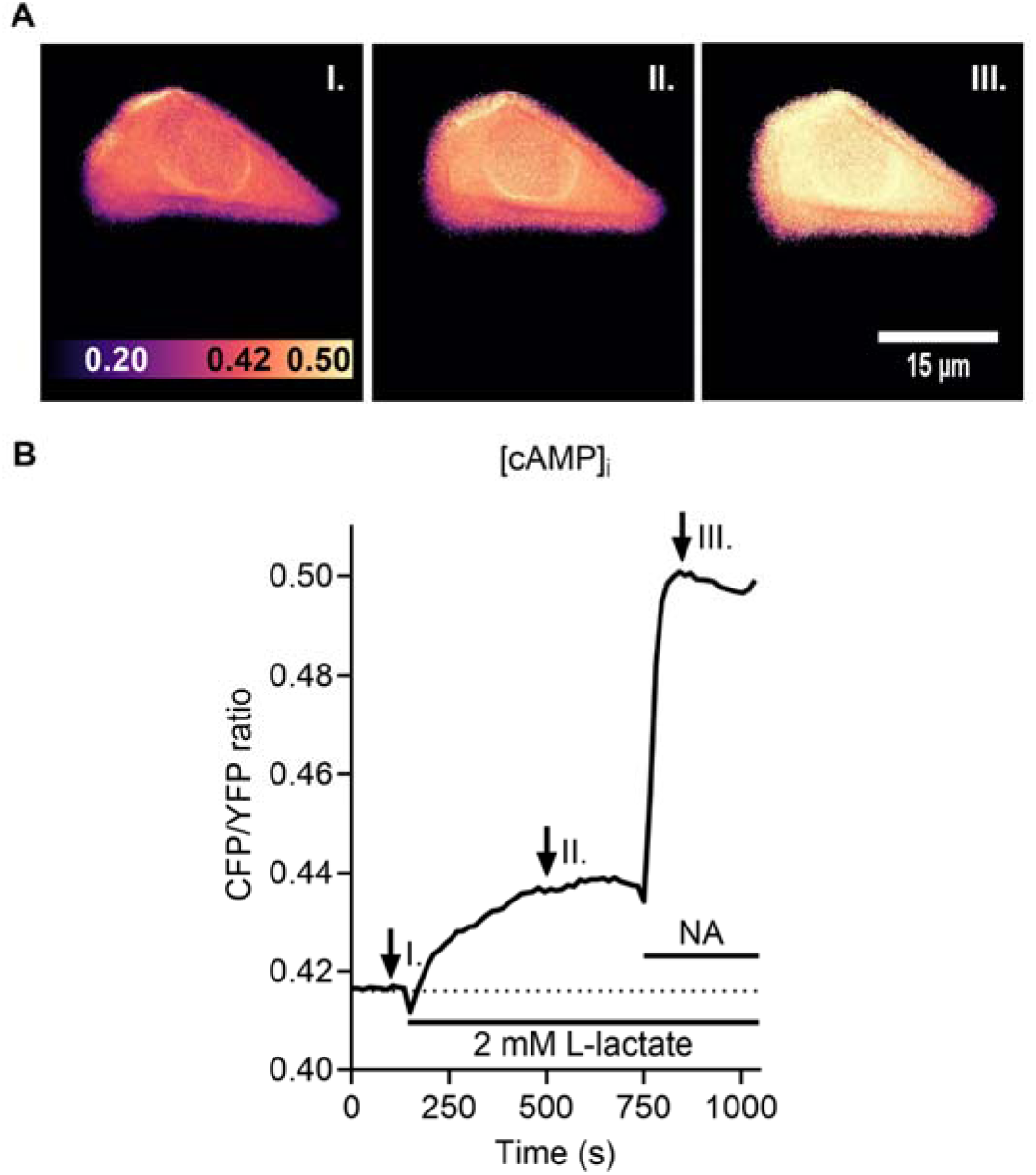
Representative single-cell [cAMP]_i_ response to extracellular L-lactate and noradrenaline (NA) in a 3T3 WT cell. (**A**) Pseudocoloured micrographs showing the CFP/YFP ratio before stimulation (I), after 2 mM L-lactate (II) and after 100 µM noradrenaline (NA; III) in a single 3T3 WT cell. Scale bar: 15 µm. (**B**) CFP/YFP ratio from the same cytoplasmic region over time. Note the rise in the signal after application of 2 mM L-lactate and the larger response to NA. The dotted horizontal line indicates the baseline CFP/YFP ratio; horizontal bars indicate L-lactate and NA application. Arrows indicate the time points shown in (**A**). For clarity, every third data point is shown. CFP, cyan fluorescent protein; WT, wild-type; YFP, yellow fluorescent protein.

Time-dependent changes in the FRET [cAMP]_i_ measurements are plotted from 0 to 730 s, with every third point shown. Responses to the application of 8535n (1 µM, Fig. 4) were quantified as AUC relative to baseline over 600 s after stimulation. Mean [cAMP]_i_ AUC ± SEM values (% × s) were: WT vehicle, 10.62 ± 2.23 (*n* = 8) and WT 8535n, 14.28 ± 1.96 (*n* = 13). Median AUC values, with 5th and 95th percentiles (% × s), were: WT vehicle, 11.48 (1.33, 17.20) and WT 8535n, 14.43 (3.40, 21.28). The difference between the groups was not statistically significant (Mann–Whitney rank-sum test, *P* = 0.262, *U* = 36). WT cells did not respond to the 8535n stimulus by increasing [cAMP]_i_ compared with vehicle-treated cells.

**Fig. 4.**
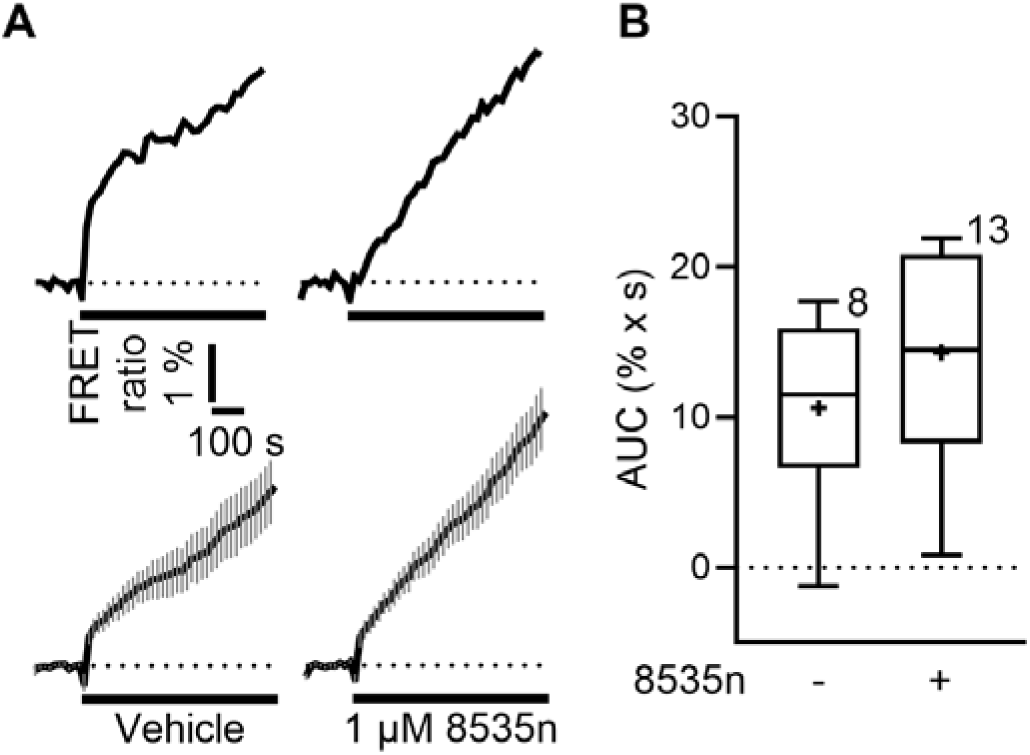
8535n does not produce an agonist-specific increase in [cAMP]_i_ in 3T3 WT cells under these experimental conditions. (**A**) Representative traces (upper panels) and averaged traces (mean ± SEM; lower panels) of the normalized CFP/YFP FRET ratio after vehicle (ECS; left) or 1 µM 8535n (right) in 3T3 WT cells. Dotted horizontal lines indicate baseline (0%), and bold horizontal bars indicate stimulus application. FRET recordings are plotted from 0 to 730 s, with every third point shown. (**B**) Box-and-whisker plots of AUC for normalized [cAMP]_i_ responses to vehicle and 8535n. Boxes show the interquartile range, horizontal lines indicate medians, whiskers show the 5th and 95th percentiles, and + indicates the mean. Numbers near boxes indicate independently recorded single-cell observations from at least three cell passages. No statistical difference, assessed by the Mann–Whitney rank-sum test, was noted (*P* = 0.262). AUC, area under the curve; CFP, cyan fluorescent protein; ECS, extracellular solution; FRET, Förster resonance energy transfer; WT, wild-type; YFP, yellow fluorescent protein.

These results show that 8535n induces a GPR27-dependent increase in intracellular Ca^2+^ but does not elicit a significant increase in [cAMP]_i_ in 3T3 WT cells.

### Extracellular L-lactate increases [Ca^2+^]_i_ and [cAMP]_i_ in 3T3 cells

We next examined how extracellular L-lactate affects the second messengers Ca^2+^ and cAMP using genetically encoded FRET nanosensors as in previous experiments (Figs. 2 and 4): D3cpv for [Ca^2+^]_i_ and Epac1-camps for [cAMP]_i_. Cells (WT, GPR27KO and rescued cells (GPR27KO + pFLAG27)) were stimulated with 2 mM extracellular L-lactate, and vehicle controls received an equal volume of ECS to match solution exchange and potential mechanical perturbation (Figs. 5 and 6).

**Fig. 5.**
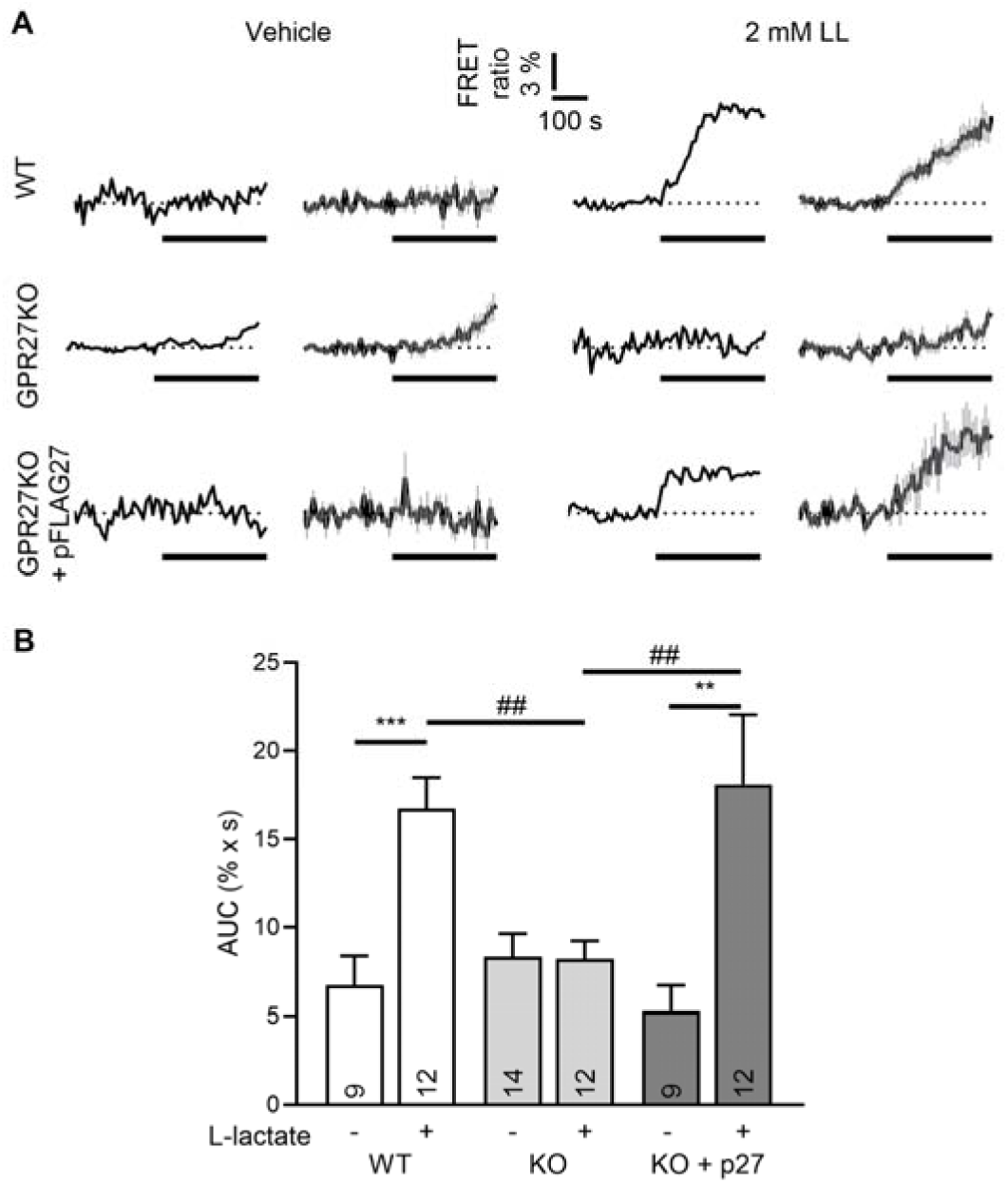
L-Lactate (2 mM) increases [Ca^2+^]_i_ in 3T3 WT cells, and the response is rescued by re-expression of GPR27 in GPR27KO cells. (**A**) Representative traces (first and third columns) and averaged traces (mean ± SEM; second and fourth columns) of the normalized YFP/CFP FRET ratio in 3T3 WT, 3T3 GPR27KO and GPR27KO cells transiently re-expressing FLAG-tagged GPR27 (GPR27KO + p27). Cells were treated with vehicle (ECS; left) or 2 mM L-lactate (LL; right). Dotted horizontal lines indicate the normalized baseline (0%), and bold horizontal bars indicate stimulus application. FRET recordings are plotted from 60 to 600 s, with every third point shown. (**B**) Mean AUC ± SEM of the normalized [Ca^2+^]_i_ response after vehicle or L-lactate. Numbers in the bars indicate independently recorded single-cell observations from at least three cell passages. Statistical significance was assessed by Student’s *t* test for WT (\*\*\**P* ≤ 0.001) and for GPR27KO (*P* = 0.936), the Mann–Whitney rank-sum test for GPR27KO + p27 (\*\**P* ≤ 0.01), one-way ANOVA for vehicle (*P* = 0.330) and Kruskal-Wallis test followed by Tukey’s post hoc test for L-lactate-treated groups (^##^*P* ≤ 0.01). AUC, area under the curve; CFP, cyan fluorescent protein; ECS, extracellular solution; FRET, Förster resonance energy transfer; KO, knockout; WT, wild-type; YFP, yellow fluorescent protein.

**Fig. 6.**
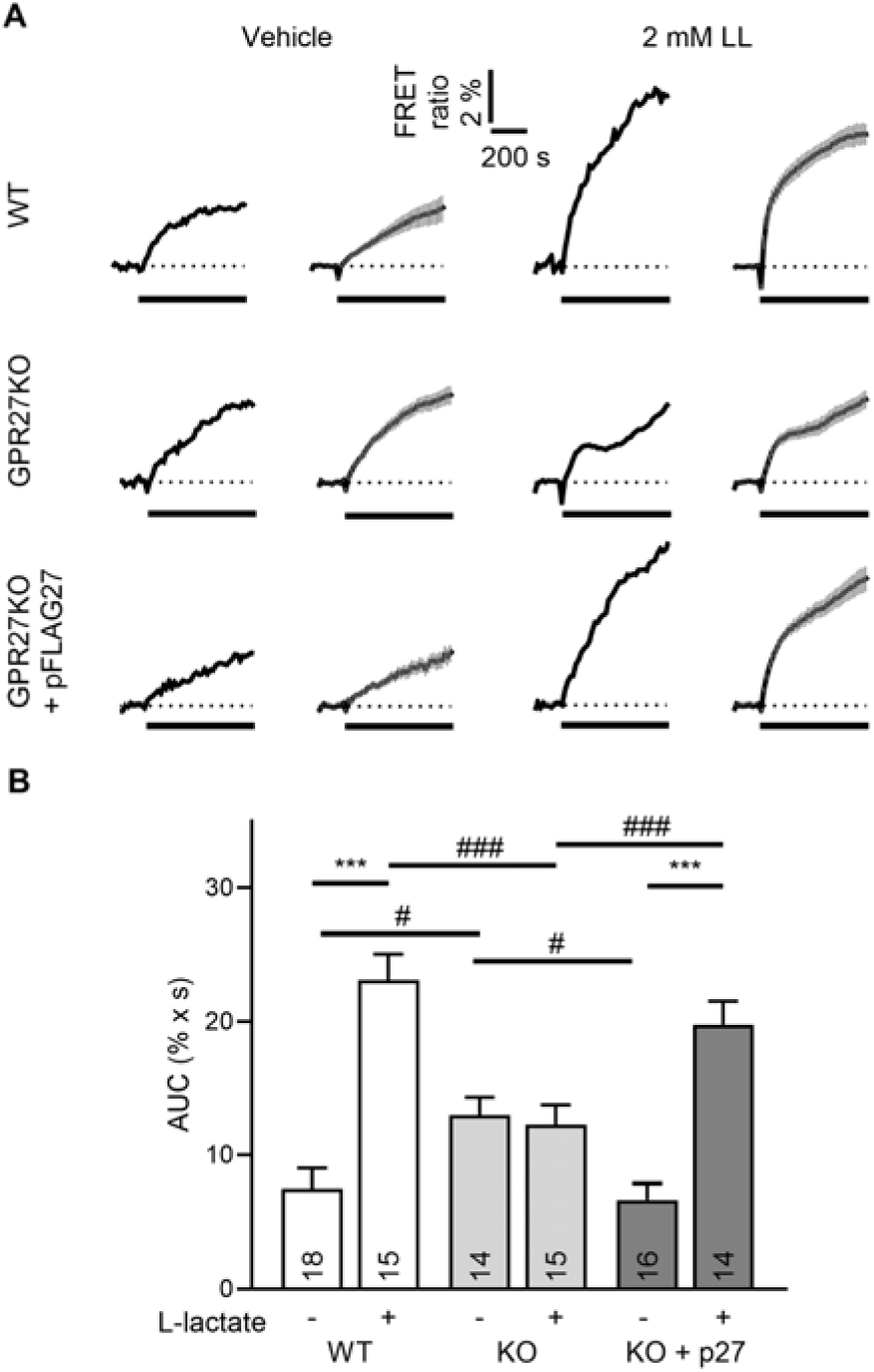
L-Lactate (2 mM) increases [cAMP]_i_ in 3T3 WT cells, and the response is rescued by re-expression of GPR27 in GPR27KO cells. (**A**) Representative traces (first and third columns) and averaged traces (mean ± SEM; second and fourth columns) of the normalized CFP/YFP FRET ratio in 3T3 WT, 3T3 GPR27KO and GPR27KO cells transiently re-expressing FLAG-tagged GPR27 (GPR27KO + p27). Cells were treated with vehicle (ECS; left) or 2 mM L-lactate (LL; right). Dotted horizontal lines indicate baseline (0%), and bold horizontal bars indicate stimulus application. FRET recordings are plotted from 0 to 730 s, with every third point shown. (**B**) Mean AUC ± SEM of normalized [cAMP]_i_ responses to vehicle or L-lactate. Numbers in bars indicate independently recorded single-cell observations from at least three cell passages. Statistical significance was assessed by the Mann–Whitney rank-sum test for WT (\*\*\**P* ≤ 0.001), Student’s *t* test for GPR27KO (*P* = 0.725), and the Mann–Whitney rank-sum test for GPR27KO + p27 (***p ≤ 0.001). Kruskal-Wallis’s test, followed by Dunn’s post hoc test, was used to compare within the L-lactate group (^###^*P* ≤ 0.001) and the vehicle group (^#^*P* ≤ 0.05). AUC, area under the curve; CFP, cyan fluorescent protein; ECS, extracellular solution; FRET, Förster resonance energy transfer; KO, knockout; WT, wild-type; YFP, yellow fluorescent protein.

For clarity, FRET recordings of increased [Ca^2+^]_i_ are shown from 60 to 600 s, with every third data point plotted (Fig. 5). Responses were quantified as the AUC relative to baseline during the 300 s after stimulation. Mean AUC ± SEM values (% × s) were: WT vehicle, 6.73 ± 1.66 (*n* = 9); WT L-lactate, 16.72 ± 1.76 (*n* = 12); GPR27KO vehicle, 8.34 ± 1.30 (*n* = 14); GPR27KO L-lactate, 8.20 ± 1.04 (*n* = 12); GPR27KO + pFLAG27 vehicle, 5.29 ± 1.44 (*n* = 9); and GPR27KO + pFLAG27 L-lactate, 18.08 ± 3.97 (*n* = 12). L-Lactate significantly increased [Ca^2+^]_i_ in WT cells (Student’s *t* test, \*\*\**P* ≤ 0.001, *t* = −4.0) and GPR27-rescued cells (Mann–Whitney test, \*\**P* ≤ 0.01, *U* = 12), but not in GPR27KO cells (Student’s *t* test, *P* = 0.936, *t* = 0.08). Vehicle responses did not differ among genotypes (one-way ANOVA, *P* = 0.330, *F* = 1.2), whereas L-lactate responses differed significantly (Kruskal-Wallis test with Tukey’s post hoc analysis, ^##^*P ≤* 0.01, *H* = 11.5; *q*_WT vs. KO_ = 4.5, *q*_WT vs. KO + p27_ = 0.9, *q*_KO + p27 vs. KO_ = 3.6). Thus, 2 mM L-lactate increased [Ca^2+^]_i_ in WT and GPR27-rescued cells, but not in GPR27KO cells, supporting a GPR27-dependent Ca^2+^ response.

For clarity, FRET recordings of increased [cAMP]_i_ are shown from 0 to 730 s, with every third data point plotted (Fig. 6). Responses were quantified as the AUC relative to baseline during the 600 s after stimulation. Mean AUC ± SEM values (% × s) were: WT vehicle, 7.50 ± 1.53 (*n* = 18); WT L-lactate, 23.08 ± 1.97 (*n* = 15); GPR27KO vehicle, 12.97 ± 1.36 (*n* = 14); GPR27KO L-lactate, 12.25 ± 1.51 (*n* = 15); GPR27KO + pFLAG27 vehicle, 6.68 ± 1.22 (*n* = 16); and GPR27KO + pFLAG27 L-lactate, 19.77 ± 1.75 (*n* = 14). L-Lactate significantly increased [cAMP]_i_ in WT and GPR27-rescued cells (Mann–Whitney test, \*\*\**P ≤* 0.001; *U* = 12 and 6, respectively), but not in GPR27KO cells (Student’s *t* test, *P* = 0.725, *t* = 0.4). Kruskal-Wallis test followed by Dunn’s post hoc test revealed significant genotype-dependent differences in both the L-lactate group (^###^*P ≤* 0.001, *H* = 15.8; *Q*_WT vs. KO_ = 3.9, *Q*_WT vs. KO + p27_ = 1.2, *Q*_KO + p27 vs. KO_ = 2.6) and the vehicle group (^#^*P ≤* 0.05, *H* = 8.2; *Q*_WT vs. KO_ = 2.5, *Q*_WT vs. KO + p27_ = 0.2, *Q*_KO + p27 vs. KO_ = 2.6). These findings indicate that GPR27 mediates the L-lactate-associated increase in [cAMP]_i_. However, the vehicle AUC in GPR27KO cells was approximately 1.8-fold higher than in WT and rescued cells, suggesting increased sensitivity to solution exchange or putative mechanical stimulation.

These results show that 2 mM L-lactate invokes a GPR27-dependent increase in intracellular Ca^2+^ and cAMP in 3T3 WT cells. Therefore, the current data support a role for GPR27 in lactate-associated Ca^2+^ and cAMP signalling.

### Basal values of [cAMP]_i_ and [Ca^2+^]_i_ are similar in 3T3 WT and GPR27KO cells

We compared baseline FRET ratios in cells expressing D3cpv or Epac1-camps before adding any stimulus to determine whether GPR27 knockout affects basal intracellular Ca^2+^ or cAMP readouts. In this context, we inspected baseline YFP/CFP and baseline CFP/YFP FRET ratios as proxies for [Ca^2+^]_i_ and [cAMP]_i,_ respectively (Fig. 7).

**Fig. 7.**
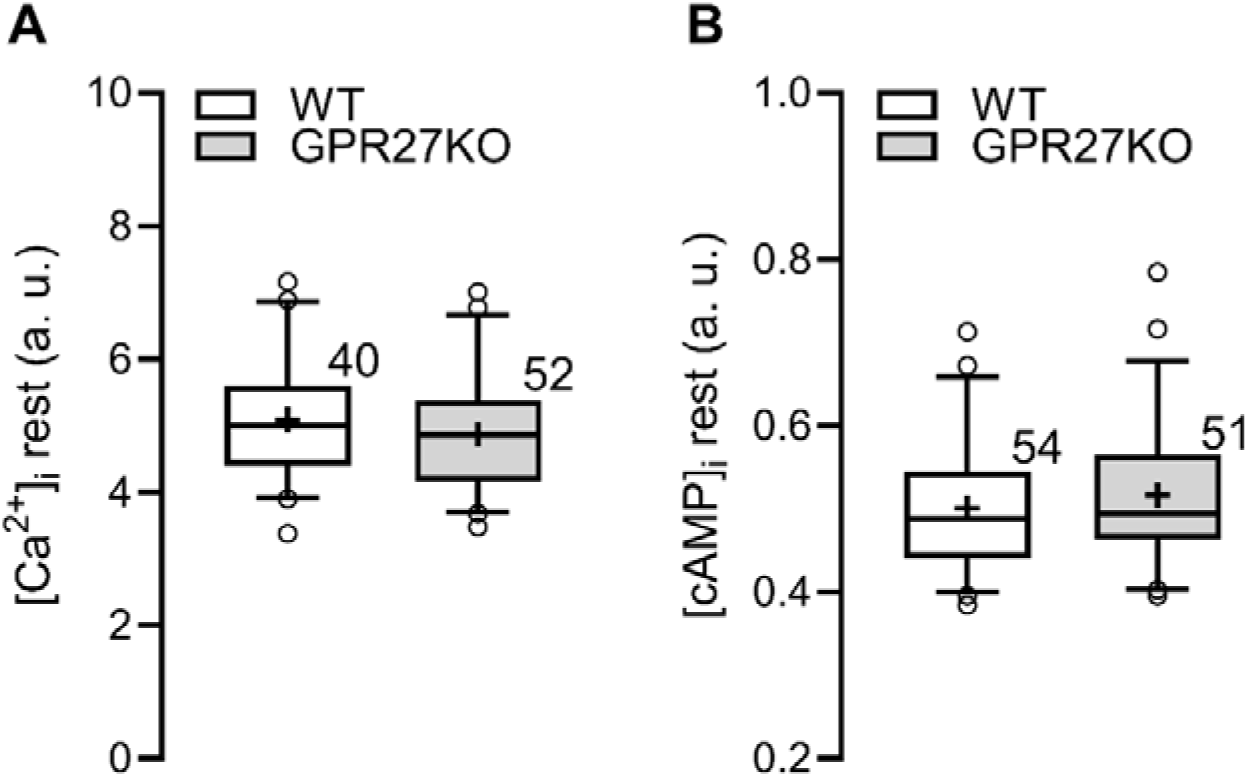
Basal Ca^2+^ and cAMP FRET ratios are similar in 3T3 WT and GPR27KO cells. (**A**) Box-and-whisker plots of baseline YFP/CFP FRET ratios, used as a basal [Ca^2+^]_i_ readout, in 3T3 WT and 3T3 GPR27KO cells. (**B**) Box-and-whisker plots of baseline CFP/YFP FRET ratios, used as a resting [cAMP]_i_ readout, in 3T3 WT and GPR27KO cells. Boxes show the interquartile range, horizontal lines indicate medians, whiskers show the 5th and 95th percentiles, and + indicates the mean. Numbers near boxes indicate independently recorded single-cell observations from at least three cell passages. Statistical significance was assessed by Student’s *t* test for the Ca^2+^ readout (*P* = 0.213) and the Mann–Whitney rank-sum test for the cAMP readout (*P* = 0.338). CFP, cyan fluorescent protein; FRET, Förster resonance energy transfer; KO, knockout; WT, wild-type; YFP, yellow fluorescent protein.

Mean basal Ca^2+^ ratio values ± SEM (a.u.) were WT, 5.09 ± 0.13 (*n* = 40) and GPR27KO, 4.87 ± 0.12 (*n* = 52). Median values, with 5th and 95th percentiles (a.u.), were: WT, 5.00 (4.10, 6.48) and GPR27KO, 4.86 (3.72, 6.27); the difference was not statistically significant (Student’s *t* test, *P* = 0.213, *t* = 1.3). Mean basal cAMP ratio values ± SEM (a.u.) were: WT, 0.50 ± 0.01 (*n* = 54) and GPR27KO, 0.52 ± 0.01 (*n* = 51). Median values, with 5th and 95th percentiles (a.u.), were WT, 0.49 (0.40, 0.65) and GPR27KO, 0.49 (0.40, 0.65); the difference was not statistically significant (Mann–Whitney rank-sum test, *P* = 0.338, *U* = 1227).

These results indicate that, under the present recording conditions, GPR27 deletion does not measurably alter basal Ca^2+^ or cAMP FRET ratios in 3T3 cells.

## Discussion

In this study, we used single-cell FRET assays to study GPR27-dependent second-messenger responses in 3T3 MEF cells. Three principal findings emerged. First, the GPR27 surrogate agonist 8535n increased [Ca^2+^]_i_ in WT cells but not in GPR27KO cells; the response was restored by re-expression of GPR27. Second, 8535n did not induce a significant agonist-specific increase in [cAMP]_i_. Third, extracellular L-lactate increased [Ca^2+^]_i_ and [cAMP]_i_ in WT and GPR27-rescued cells but not in GPR27KO cells. GPR27 deletion also altered the vehicle response in [cAMP]_i_ recordings. These results implicate GPR27 in Ca^2+^ and cAMP signalling and suggest that L-lactate may act as a modulator or ligand of GPR27.

The 8535n-induced Ca^2+^ response is consistent with previous evidence linking GPR27 to phospholipase C signalling. It was reported that GPR27 expression in HEK293T cells increased IP_1_ levels without altering cAMP and that GPR27 knockdown in pancreatic beta cells reduced IP_1_ signalling^13^. Our single-cell FRET data extend these observations by showing that pharmacological stimulation produces a GPR27-dependent Ca^2+^ response in intact 3T3 cells. Although this finding is compatible with GPR27–G_q/11_ coupling, it does not prove direct coupling. The Ca^2+^ increase could also arise through phospholipase C–IP_3_ signalling, store-operated Ca^2+^ entry, receptor crosstalk, β-arrestin-dependent scaffolding or other indirect pathways. The absence of an agonist-specific cAMP response to 8535n is also consistent with the unchanged cAMP levels reported after GPR27 expression in HEK293T cells^13^. This argues against robust GPR27–G_s_ coupling in 3T3 cells but does not exclude G_i/o_ coupling, which will have to be elucidated in future experiments.

Extracellular L-lactate produced a GPR27-dependent Ca^2+^ response that was abolished by GPR27 deletion and restored by GPR27 receptor re-expression. This provides complementary evidence that GPR27 participates in lactate-induced Ca^2+^ signalling and raises the possibility that L-lactate may be an endogenous ligand or modulator of GPR27. However, the molecular basis of this interaction remains unresolved. Lactate may act directly on GPR27 or indirectly through receptor crosstalk, transport-dependent metabolic effects or signalling pathways upstream of GPR27.

Unlike 8535n, L-lactate also increased [cAMP]_i_. This finding requires particular caution because L-lactate is an established extracellular signalling metabolite acting through HCAR1/GPR81, a G_i_-coupled receptor that generally inhibits adenylyl cyclase and lowers cAMP in adipocytes^19^. Nevertheless, increased intracellular lactate and cAMP levels have previously been observed in astrocytes, raising the possibility of cell-type-specific or non-canonical HCAR_1_ signalling^18^. In the present experiments, 2 mM extracellular L-lactate increased the Epac1-camps CFP/YFP signal in WT and GPR27-rescued cells. This response is inconsistent with canonical HCAR_1_ signalling and may instead reflect cell-specific cAMP regulation, lactate transport or metabolism, pH-dependent effects, osmotic or sodium-dependent mechanisms, receptor crosstalk or GPR27-dependent modulation of mechanosensitive cAMP production. Loss of the response in GPR27KO cells and its restoration after GPR27 re-expression demonstrate that GPR27 is required for the phenotype, but do not establish that L-lactate directly activates GPR27 through G_s_.

The increased vehicle-induced cAMP response in GPR27KO cells suggests that GPR27 deletion alters sensitivity to solution exchange or mechanical perturbation. This enhanced background response may mask or distort L-lactate-specific effects in the knockout cells and may also explain the apparent absence of an 8535n-induced cAMP response. It was reported recently that a brief pressure pulse dose-dependently increases intracellular L-lactate production in astrocytes^29^. It remains to be seen whether the vehicle-induced increase in [cAMP]_i_ observed in our work may be a mechanically driven artefact.

Basal Ca^2+^ and cAMP FRET ratios were similar in WT and GPR27KO cells, indicating that GPR27 deletion does not measurably alter basal levels of these second messengers under the present conditions. This contrasts with the previously reported increase in basal intracellular lactate in GPR27KO 3T3 cells^16^, suggesting that GPR27 may regulate cellular metabolism without producing a sustained global change in cytosolic Ca^2+^ or cAMP. Alternatively, basal effects may be confined to signalling microdomains that are not resolved by the cytosolic sensors used here.

These findings help place GPR27 within the broader context of its reported roles in insulin-promoter regulation, glucose homeostasis, lipid metabolism, lactate homeostasis and the pathophysiology of neurodegeneration and cancer. The disease groups mentioned exhibit metabolic changes. A hypoenergetic brain state is observed clinically in Alzheimer disease^30^. In Parkinson disease, metabolic impairment is complex; however, FDG-PET hypometabolism studies identified consistent functional brain abnormalities^31^. A metabolic shift is often observed in cancer cells^17^. These changes may be associated with GPR27.

Because Ca^2+^ and cAMP regulate transcription, secretion, metabolism, proliferation and ERK/MAPK activity, the L-lactate-induced increase in [Ca^2+^]_i_ and [cAMP]_i_, requiring GPR27, may contribute to the complexity of metabolic changes in neurodegeneration and cancer. However, the relationship between acute signalling in 3T3 MEF cells and the longer-term phenotypes observed in other cell types remains to be established.

Overall, our data support a model in which GPR27 regulates Ca^2+^ signalling in 3T3 MEF cells and contributes to Ca^2+^ and cAMP responses elicited by extracellular L-lactate. GPR27 may also modulate cellular sensitivity to mechanical stimulation. Although the findings are consistent with L-lactate acting as an endogenous GPR27 ligand or modulator, direct ligand–receptor binding and the underlying G-protein coupling mechanisms remain to be demonstrated.

## Conclusion

In 3T3 MEF cells, the GPR27 surrogate agonist 8535n induced a GPR27-dependent increase in [Ca²⁺]_i_ without significantly affecting [cAMP]_i_. L-Lactate similarly increased [Ca²⁺]_i_ in WT and GPR27-rescued cells, but not in GPR27KO cells, and enhanced [cAMP]_i_ in WT and GPR27-rescued cells. These findings suggest that GPR27 mediates L-lactate-associated Ca²⁺ and cAMP signalling.

## Methods

### Cell culture and plasmid transfection

3T3 MEF cells were kindly provided by Dr K. Chylinski (Vienna BioCenter, University of Vienna, Vienna, Austria). Experiments were conducted with 3T3 WT cells and CRISPR-Cas9-edited 3T3 GPR27KO cells (IE1 clone carrying a homozygous 31-bp deletion), as previously described^16^. GPR27KO 3T3 cells were sequenced to validate that the *Gpr27* gene was knocked out from the 3T3 cells (Supplementary Fig. S1). Cultures were maintained in high-glucose Dulbecco’s modified Eagle’s medium (Sigma-Aldrich, St. Louis, MO, USA, cat. no. D6546) supplemented with 10% fetal bovine serum (Sigma-Aldrich, cat. no. F-7524), 5 mM L-glutamine (Sigma-Aldrich, cat. no. G-3126) and 25 µg/mL penicillin-streptomycin (Sigma-Aldrich, cat. no. P-0781) in a humidified incubator (95% air, 5% CO_2_, 37°C). Cells were subcultured every 2 days when they reached 70%–80% confluence. Each experimental series included cells from at least three different passages, ranging from 2 to 10. For the experiments, cells were detached using trypsin-(ethane-1,2-diamine) tetraacetic acid (Sigma-Aldrich, cat. no. T-3924) and seeded onto poly-L-lysine-coated coverslips (Sigma-Aldrich, cat. no. P-1524). Cultured cells were maintained in complete growth medium until experimentation.

At least 24 h before imaging, cells were transfected with the genetically encoded FRET-based nanosensor pcDNA-D3cpv (Addgene, Watertown, MA, USA; plasmid cat. no. 36323), Epac1-camps (a generous gift from Martin J. Lohse, Institute of Pharmacology and Toxicology, University of Würzburg, Germany) or ssFlagGPR27 (a generous gift from Dr Julien Hanson, University of Liège, Belgium). The transfection mixture (FuGene 6 Transfection Reagent; Promega, Madison, WI, USA, cat. no. E-2692; plus 0.5 µg plasmid DNA) was prepared in antibiotic- and serum-free medium, applied for 2 h at 37°C, and subsequently replaced with standard culture medium.

### Experimental extracellular solutions

The ECS contained 135.3 mM NaCl (Sigma-Aldrich, cat. no. S-7653), 5 mM KCl (Sigma-Aldrich, cat. no. P3911), 10 mM HEPES (4-(2-hydroxyethyl)-1-piperazineethanesulfonic acid; Sigma-Aldrich, cat. no. H-3375), 0.5 mM NaH_2_PO_4_·H_2_O (Sigma-Aldrich, cat. no. S0751), 5 mM NaHCO_3_ (Sigma-Aldrich, cat. no. S-5761), 2 mM MgCl_2_ (Sigma-Aldrich, cat. no. M-8266), 1.8 mM CaCl_2_ (Sigma-Aldrich, cat. no. 21115) and 3 mM D-glucose (Sigma-Aldrich, cat. no. G8279). The pH was adjusted to 7.2 with NaOH (Merck, Darmstadt, Germany, cat. no. 1064981000) using a SevenEasy S20 pH meter (Mettler Toledo, Columbus, OH, USA). Osmolarity was maintained at 290–310 mOsm and measured with a freezing-point osmometer (Osmomat 030; Gonotec GmbH, Berlin, Germany).

The surrogate GPR27 agonist 8535n (N-[4-(anilinosulphonyl)phenyl]-2,4-dichlorobenzamide)^1, 24, 32^ was used (ChemBridge Corporation, San Diego, CA, USA, cat. no. SC-5128535) to monitor changes in intracellular second messengers after stimulation of GPR27. Stock solutions were prepared at 10 mM in dimethyl sulfoxide (DMSO; Sigma-Aldrich, cat. no. D8418) and diluted into ECS to obtain a final working concentration of 1 µM. The final DMSO concentration applied to cells did not exceed 0.01%.

Changes in intracellular second messengers after stimulation with L-lactate were monitored by modifying the composition of the ECS to include L-lactate while maintaining osmolarity within the same range (290–310 mOsm, pH 7.2). Specifically, the solution contained 95.3 mM NaCl, 5 mM KCl, 10 mM HEPES, 0.5 mM NaH_2_PO_4_·H_2_O, 5 mM NaHCO_3_, 2 mM MgCl_2_, 1.8 mM CaCl_2_ and 3 mM D-glucose, supplemented with sodium L-lactate (Sigma-Aldrich, cat. no. 71718) to obtain a 40 mM stock solution. This stock solution was then diluted with ECS as previously described to achieve a final concentration of 2 mM L-lactate.

Ionomycin (10 µM; Sigma-Aldrich, cat. no. I0634) diluted in ECS was used as a positive control for Ca^2+^ measurements. NA (100 µM; Sigma-Aldrich, cat. no. A7256) diluted in ECS served as the positive control for cAMP measurements.

### FRET measurements and data analysis

FRET imaging was used to monitor intracellular Ca^2+^ and cAMP in real time in single 3T3 cells. Transfected cells were incubated for 30 min in standard ECS and mounted in a custom recording chamber containing 200 µl of ECS. After a baseline recording of 300 s for Ca^2+^ measurements or 150 s for cAMP measurements, an equal volume (200 µl) of vehicle, surrogate agonist or L-lactate solution was added. All stimulation solutions were prepared at 2× final concentration in ECS so that the stated final concentration was reached after addition to the recording chamber. Recording continued for an additional 600 s, followed by application of 400 µl of the appropriate positive-control solution to verify sensor responsiveness: 10 µM ionomycin in Ca^2+^ measurements and 100 µM NA in cAMP measurements. The detailed cell stimulation programmes for Ca^2+^ and cAMP signal recording are shown in Figs 1 and 3.

Imaging was performed 24–36 h after transfection using a Zeiss Axio Observer.A1 fluorescence microscope (Zeiss, Oberkochen, Germany) equipped with a C-Apochromat 40×/1.3 NA oil-immersion objective (Zeiss) and an Axiocam 702 mono digital camera (Zeiss). Cells were excited at 430 nm using a Colibri 7 light-emitting diode module (Zeiss). Emitted fluorescence was separated into cyan (460 nm) and yellow (520 nm) channels by an image splitter (Photometrics DV2; Optical Insights, Tucson, AZ, USA). Images were acquired every 3 s for Ca^2+^ measurements and every 5 s for cAMP measurements, with exposure times of 80 ms and 1 s, respectively. For the Ca^2+^ sensor D3cpv, relative changes in [Ca^2+^]_i_ were calculated as changes in the YFP/CFP ratio; an increase in the ratio corresponds to an increase in [Ca^2+^]_i_. Epac1-camps reports cAMP as a decrease in FRET; therefore, cAMP data were plotted as CFP/YFP so that upward deflections consistently represent increases in [cAMP]_i_. The baseline FRET ratio for each cell was defined as the mean value during the pre-stimulus period. Changes in the FRET ratio fluorescence within a region of interest were measured by outlining the cell and subtracting the background fluorescence from both signals. Ratio traces were corrected for photobleaching-related baseline drift using a custom MATLAB routine (fitting the initial steady-state segment, extrapolating it over the recording, and subtracting it). The AUC was calculated for each normalized-to-baseline response curve from addition of stimulus to 300 s after stimulation for Ca^2+^ measurements, and from addition of stimulus to the application of the positive control for cAMP measurements (600 s).

### Statistical analysis

Unless otherwise stated, results are presented as means ± standard error of the mean (SEM). Statistical analyses were performed using SigmaPlot 11.0, Microsoft Excel and GraphPad Prism 11.0. Differences were considered statistically significant at *P ≤* 0.05, *P ≤* 0.01 and *P ≤* 0.001 and are indicated in the figures by one, two or three asterisks and hashtags, respectively. In the box-and-whisker plots, boxes represent the interquartile range (IQR; middle 50% of the data), horizontal black lines indicate medians, + indicates the mean, and whiskers extend to the 5th and 95th percentiles. In the bar plots, columns show mean values and error bars indicate the SEM. For comparisons between two groups, statistical significance was assessed using a two-tailed Student’s *t* test when the data met the assumptions of normality and equal variance, as verified by Shapiro–Wilk and Levene tests. When these assumptions were violated, the Mann–Whitney rank-sum test was applied. For comparisons involving more than two groups, one-way ANOVA was used, followed by the indicated post hoc tests. *n* denotes independently recorded single-cell observations. Cells were obtained from at least three cell passages. *t*, the *t* statistic used in Student’s *t* test; U, the *U* statistic used in the Mann–Whitney rank-sum test; *F*, the *F* statistic used in one-way ANOVA or the Friedman test; *H*, the *H* statistic used in the Kruskal-Wallis test; *Q*, *t* and *q*, the test statistics used in post hoc analyses. Sample size was not predetermined using statistical methods, and blinding, randomization, and outlier tests were not performed.

## Abbreviations

8535n: N-[4-(anilinosulphonyl)phenyl]-2,4-dichlorobenzamide
ANOVA: analysis of variance
AUC: area under the curve
[Ca^2+^]_i_: intracellular calcium
[cAMP]_i_: intracellular cyclic adenosine monophosphate
CFP: cyan fluorescence protein
DMSO: dimethyl sulfoxide
ECS: extracellular solution
ERK: extracellular signal-regulated kinase
FDG-PET: fluorodeoxyglucose-positron emission tomography
FRET: Förster resonance energy transfer
GPCR: G-protein coupled receptor
HCAR1: hydroxycarboxylic acid receptor 1
HEK293T: human embryonic kidney cells that contain the SV40 large T antigen
HEPES: 4-(2-hydroxyethyl)-1-piperazineethanesulfonic acid
IONO: ionomycin
IP_1_: inositol monophosphate
IP_3_: inositol 1,4,5-trisphosphate
IQR: interquartile range
[lactate]_i_: intracellular L-lactate concentration
MAPK: mitogen-activated protein kinase
MEF: murine embryonic fibroblast
NA: noradrenaline
oGPCR: orphan G-protein coupled receptor
SEM: standard error of the mean
SREB: superconserved receptors expressed in the brain
WT: wild-type
YFP: yellow fluorescent protein

## Author contributions

Conceptualization: E.S., H.H.C. and R.Z.; Methodology: D.K., M.K., H.H.C. and R.Z.; Formal analysis: D.K. and T.M.Z.; Investigation: E.S.; Writing – original draft: D.K.; writing – review & editing: D.K., E.S., T.M.Z., M.K., H.H.C. and R.Z.; Visualization: D.K.; Supervision: M.K., H.H.C. and R.Z.; Project administration: R.Z.; Funding acquisition: M.K., H.H.C. and R.Z.

## Lead contact

Requests for further information and resources should be directed to and will be fulfilled by the lead contact, Robert Zorec.

## Data and code availability

Data supporting the findings of this study and original code are available from the lead contact upon reasonable request.

## Materials availability

This study did not generate new unique reagents.

---

**Supplementary Fig. S1.**
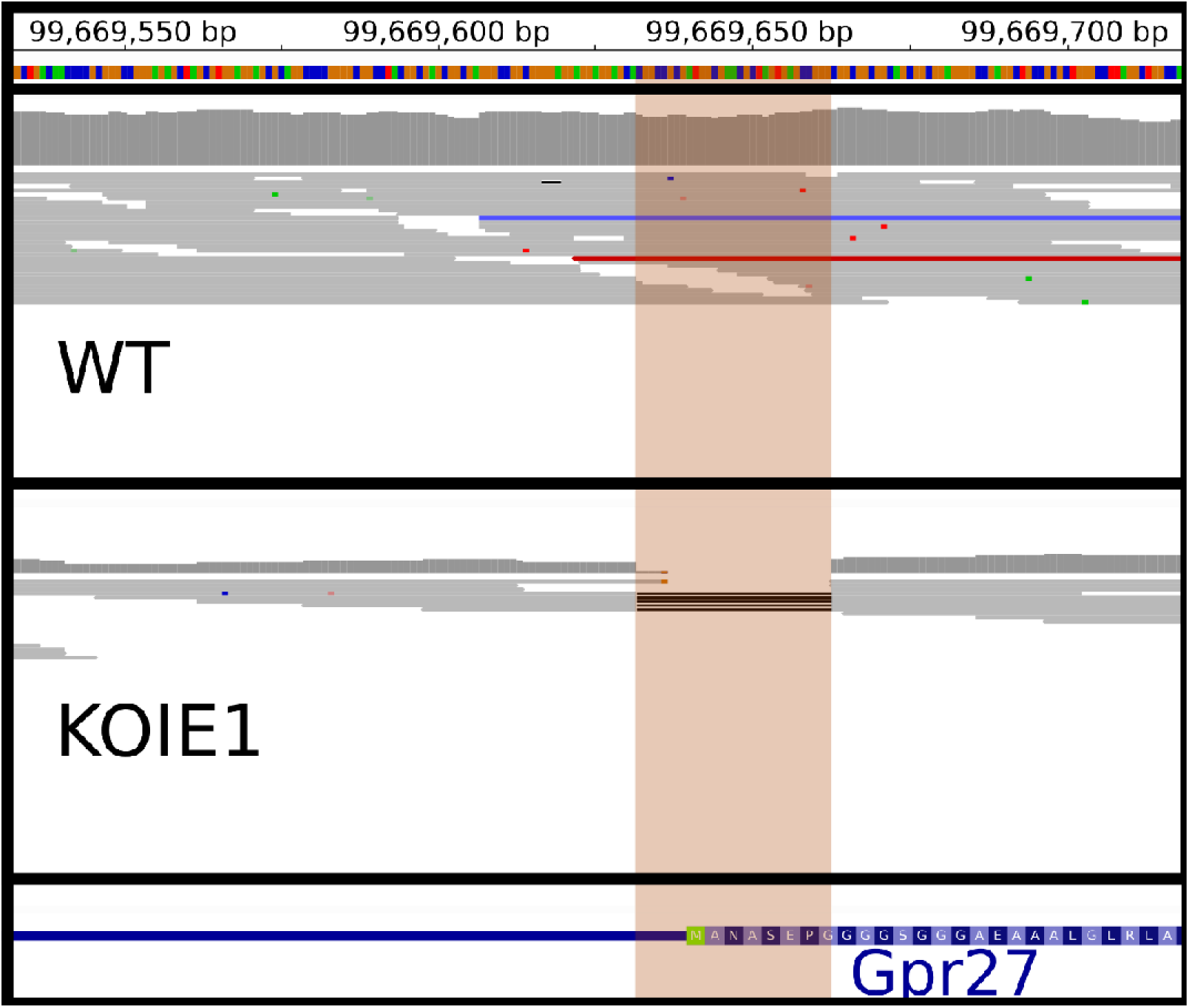
Knockout of Gpr27 in KOIE1 3T3 was achieved by deletion of the Gpr27 start codon. The top pane shows the colour-coded nucleic sequence (A, green; T, red; C, blue; A, yellow) with numeric chr6 positions in the mm39 mouse reference genome; the bottom pane indicates the gene location (blue line) and its coding sequence (single letter amino acid codes; M in green); positions match the indicated chromosome 6 positions from the top pane. Top- and bottom-middle panes indicate sequencing coverage at the chromosome 6 positions indicated in the top pane; each of these two panes indicates numeric coverage depth values at each chromosomal position at the top; below are read stacking diagrams, which show individual read placements; reads are coloured according to insert size deviation (grey, in normal range; blue, insert size lower than expected; red, insert size greater than expected); deletion of reference bases in the read is marked with black lines. The top- and bottom-middle panes correspond to genome sequencing of WT and KOIE1 3T3 cells. In the mm39 mouse genome reference, the Gpr27 coding sequence is located on chr6 at 99,669,640–99,670,776 bp. The brown transparent band indicates the position of the deleted part of the sequence in KOIE1 chr6:g.99669632_99669663del. Comparing the span of the Gpr27 coding sequence and the deletion location (bottom and bottom-middle panes) shows that Gpr27 was knocked out by deleting the immediate region surrounding its start codon. KO, knockout; WT, wild-type.

